# c-di-GMP modulates ribosome assembly by inhibiting rRNA methylation

**DOI:** 10.1101/2024.06.05.597503

**Authors:** Siqi Yu, Zheyao Hu, Xiaoting Xu, Xiaoran Liang, Jiayi Shen, Min Liu, Mingxi Lin, Hong Chen, Jordi Marti, Sheng-ce Tao, Zhaowei Xu

**Author notes:** These authors contributed equally to this study (Siqi Yu, Zheyao Hu, Xiaoting Xu). Corresponding authors. (Zhaowei Xu), (Sheng-ce Tao), (Jordi Marti).

## Abstract

Cyclic diguanosine monophosphate (c-di-GMP) is a ubiquitous bacterial secondary messenger, with diverse functions, many of which are yet to be uncovered. Stemming from an *Escherichia coli* proteome microarray, we found that c-di-GMP bound to 23S rRNA methyltransferases (RlmI and RlmE). rRNA methylation assays showed that c-di-GMP inhibits RlmI activity, thereby modulating ribosome assembly. Based on molecular dynamic simulation and mutagenesis studies, we found that c-di-GMP binds to RlmI at residues R64, R103, G114, and K201. Structural simulation revealed that c-di-GMP quenches RlmI activity by inducing the closure of the catalytic pocket. Furthermore, we revealed that c-di-GMP promotes antibiotic tolerance by regulating RlmI activity, which played a role in antibiotic-resistant strains. Finally, the binding and methylation assays showed that the effect of c-di-GMP on RlmI is conserved, at least in various pathogenic bacteria. This study discovered an unexpected functional role of c-di-GMP in regulating ribosome assembly by inhibiting rRNA methylases. This study identified an unexpected but crucial member among the c-di-GMP effectors.

**Highlights:**

1. c-di-GMP regulates ribosome assembly in *Escherichia coli*.
2. c-di-GMP inhibits rRNA methylation activity of RlmI by inducing catalytic pocket closure.
3. c-di-GMP promotes antibiotic resistance by regulating ribosome assembly.

**Graphical Abstract:** 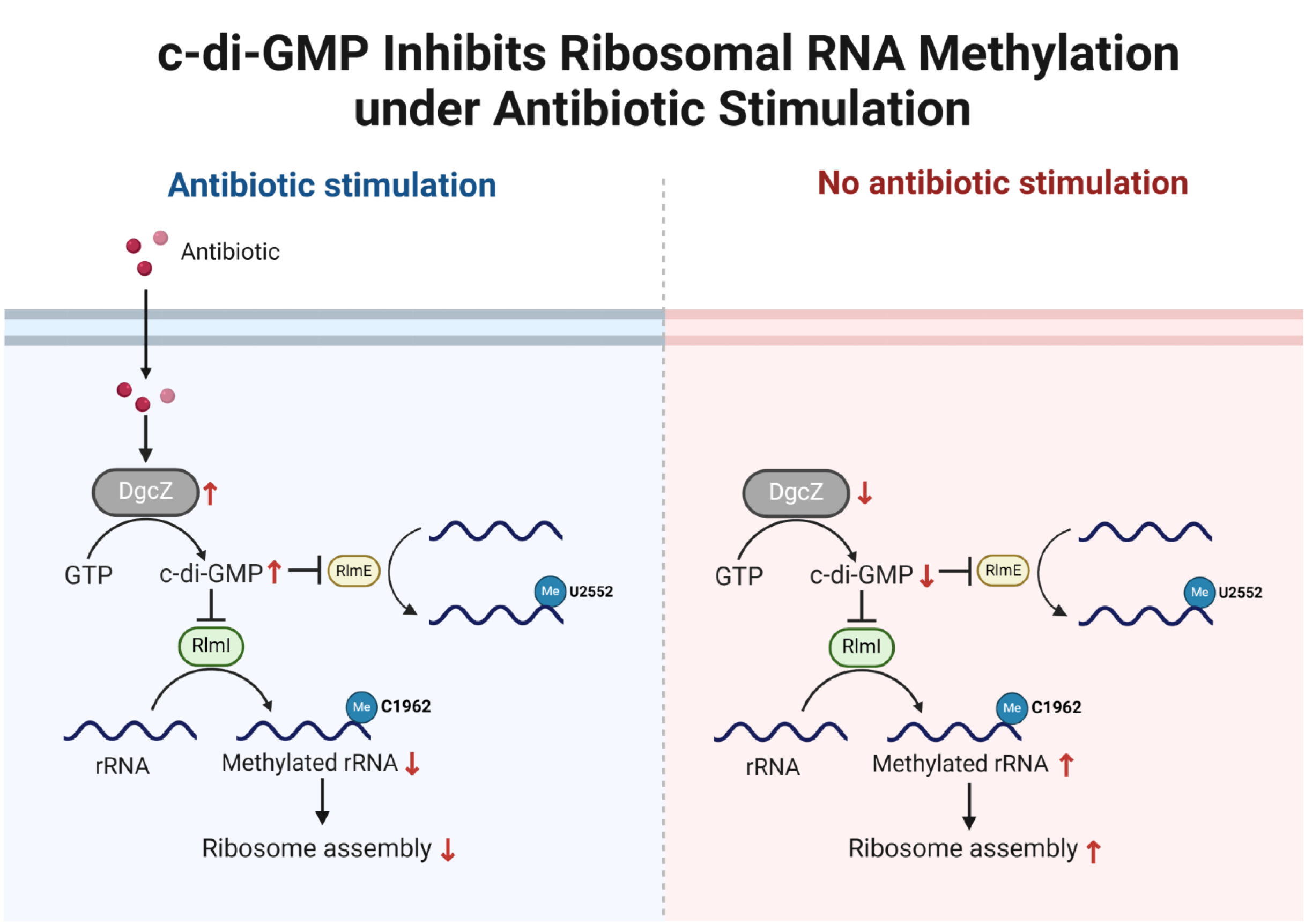

## Introduction

Cyclic diguanosine monophosphate (c-di-GMP) was first identified in *Gluconacetobacter xylinus,* where it regulated cellulose synthesis^1^. Subsequent research revealed that c-di-GMP plays a role in a wide range of bacterial biological processes, such as motility, virulence, and host-microbe symbiosis ^2–5^. In a previous study, we globally screened c-di-GMP binding proteins by applying an *Escherichia coli* proteome microarray and revealed the interplay loop between c-di-GMP and protein acetylation^6^. Interestingly, the microarray assay also showed that 23S rRNA methyltransferases RlmI and RlmE act as c-di-GMP binding proteins. This suggests a functional link between c-di-GMP and ribosome assembly.

The ribosome assembly involves the processing and folding of rRNA and its concomitant assembly with ribosomal proteins. As a part of rRNA processing, rRNA methylation participates in the regulation of ribosome assembly. For example, the inactivation of RlmE is associated with a large subunit assembly defect ^7^, and RsmA fulfills quality control requirements in the last stages of small subunit assembly ^8^. Overall, 23 ribosomal RNA methylases exist in *E. coli*, most of which have unresolved physiological functions. Consequently, studying the function and regulatory factors of rRNA methyltransferase is crucial for understanding the mechanism of ribosome assembly.

Ribosome biogenesis is a fundamental cellular process, equipping cells with the molecular factories for cellular protein production. Inhibiting ribosome assembly is considered an essential source of new targets for drug development^9,10^. Therefore, studying the relationship between ribosome assembly and bacterial resistance is crucial for designing antibiotics targeting ribosome assembly pathways. In Gram-positive bacteria, (p)ppGpp negatively impacts ribosome assembly by inhibiting GTPase activity, influencing growth and antibiotic tolerance^11^. However, in Gram-negative bacteria, the influence of (p)ppGpp on antibiotic tolerance mainly affects nucleotide and amino acid synthesis^12,13^. The regulatory relationship between ribosome assembly and antibiotic tolerance in Gram-negative bacteria is yet to be understood.

rRNA methylation serves as a significant mechanism for bacterial resistance against ribosome-targeting antibiotics. Two clinically relevant examples include 16S and 23S rRNA methyltransferases, which confer resistance by modifying conserved rRNA residues in site A or PTC, respectively. This modification makes bacteria insensitive to aminoglycosides and streptogramin B ^14^. For instance, aminoglycoside resistance in *E. coli* is conferred by methylation of the G1405 and A1408 residues in the 16S rRNA by RsmF ^15^ and NpmA ^16^, respectively. The Erm macrolide-resistance methyltransferase family has been widely identified in Gram-positive bacteria ^17^ and continued to spread and mutate globally ^18,19^. The chloramphenicol-florfenicol resistance (*cfr*) SAM methyltransferase family shares ancestry with the housekeeping RlmN methyltransferases. These methyltransferases incorporate methylation at A2503 in 23S rRNA and A37 in tRNAs ^20,21^. However, the upstream regulatory factors and downstream mechanisms of rRNA methylation in the context of antibiotic resistance remain unclear.

In this study, we demonstrate that c-di-GMP binds to four 23S rRNA methyltransferases and that RlmI is the main effector of c-di-GMP in regulating ribosome assembly. Structural analysis revealed that c-di-GMP binds to RlmI at the R64, R103, G114, and K201 residues and induces the closure of the catalytic pocket of RlmI. We further show that c-di-GMP regulates ribosomal assembly to promote antibiotic resistance by inhibiting RlmI activity. Finally, sequence comparison of RlmI orthologues among bacteria indicated that some important human pathogens are conserved in the c-di-GMP-based rRNA regulation mechanism.

## Material & Methods

### E. coli strains and plasmids

*E. coli* BW25113 as reference strain was used in this study, and pCA24N and pGEX4T-1 plasmids were used to overexpress methyltransferases and their mutants. Further, pET28a was used for RlmI of *S. typhimurium*, *K. pneumoniae*, and *V. cholerae*. We performed recombinant RlmI mutations using a QuikChange Site-Directed Mutagenesis Kit (#200518, Agilent Technologies, USA).

The strains described earlier were grown in Vogel-Bonner medium (0.81 mM MgSO_4_·7H_2_O, 43.8 mM K_2_HPO_4_, 10 mM C_6_H_8_O_7_·H_2_O, and 16.7 mM NaNH_4_HPO_4_·4H_2_O) with 10 mM acetate at 25°C for functional analysis. For kanamycin treatment, 3 μg/mL kanamycin was added to the Vogel-Bonner medium.

For protein purification, the strains were grown in Lysogeny broth (10 g tryptone, 5 g yeast extract, 10 g NaCl per 1 L) medium (LB medium), and then induced by 0.2 mM IPTG at 22°C for 12 h.

### Construction of endogenous DgcZ and RlmI mutants in E. coli

DgcZ and RlmI mutants were constructed using the Red-recombination system based on *E. coli* BW25113 strain ^22^, as discussed previously ^23^. For kanamycin-resistant *E. coli* strains, the *rlmI*^K201A^ mutants were constructed using the Red-recombination system with ampicillin as the screening antibiotic.

### Methyltransferase activity assay in vitro

For *in vitro* methylation assays, 3 μM recombinant enzymes (RlmI and RlmE) and c-di-GMP (at either 5 μM, 10 μM, or 20 μM) were incubated in 20 μL reaction buffer (40 mM HEPES, 100 mM NH_4_Cl, 10 mM MgCl_2_, 1 mM AdoMet, pH 7.6) at 37°C for 0.5 h. Then, 5 μM rRNA substrates were added and incubated at 37°C for 1.5 h. Reactions were stopped by heating at 95°C for 10 min and centrifugation at 10,000 x *g* for 10 min at 4°C to remove sediments.

These samples were analyzed by HPLC. Briefly, the samples were injected into a C-18 column (Alltima C18 4.6 250 mm^2^) and analyzed by reversed-phase HPLC (Shimadzu, Japan). Solution A [0.065% trifluoroacetic acid in 100% water (*v*/*v*)] and solution B [0.05% trifluoroacetic acid in 100% acetonitrile (*v*/*v*)] were used in a gradient program (0.01 min with 5% Solution B, 20 min with 65% Solution B, 20.01 min with 95% Solution B, 31 min with 95% Solution B, 31.01 min with 5% Solution B, 40 min with 5% Solution B, and stop in 40.01 min) with a flow rate of 1 mL/min. rRNA was detected at 220 nm, and the area under the curve was integrated for the relative quantification of reaction products. This assay was performed in triplicate, and the results were calculated using GraphPad Prism 6.

### MeRIP-qPCR quantification of the endogenous C1962 methylation of 23S rRNA

*E. coli* strains, such as WT, Δ*dgcZ*, *rlmI*^K201A^ and Δ*dgcZ rlmI*^K201A^ were cultivated in LB medium at 37°C overnight and transferred to Vogel-Bonner medium at a 1:1000 ratio with 10 mM acetate and 3 μg/mL kanamycin for 25°C for 24 h. Twenty OD cells were harvested for RNA extraction using an RNA extraction kit (Sangon Biotech, China). Total RNA (50 μg) was diluted in 200 μL of IP buffer (20 mM HEPES, 50 mM KCl, 1 U/μL protector RNase inhibitor, pH=7.5) and treated with a sonicator (Sonics and Materials, USA) (20 cycles on 35% power, 15 s on/off) to prepare the RNA fragments, which were divided into 50 μL as input and 150 μL for IP. Then, 2 μL of anti-m5C antibody was added to 150 μL of RNA fragments and incubated at 4°C for 4 h. 20 μL Protein A/G beads (Thermo Fisher Scientific, USA) were added to the mixture and incubated at 4°C for 1 h. The RNA was eluted with 0.1 M glycine (pH = 2) and neutralized with 1.0 M Tris (pH = 8). The eluted RNA and RNA fragments were purified by phenol‒chloroform and quantified using a NanoDrop 2000 spectrophotometer (Thermo Fisher Scientific). Real-time RT-PCR was performed using a reverse transcription kit and real-time PCR kit (Applied Biosystems, USA). The following primers were used.

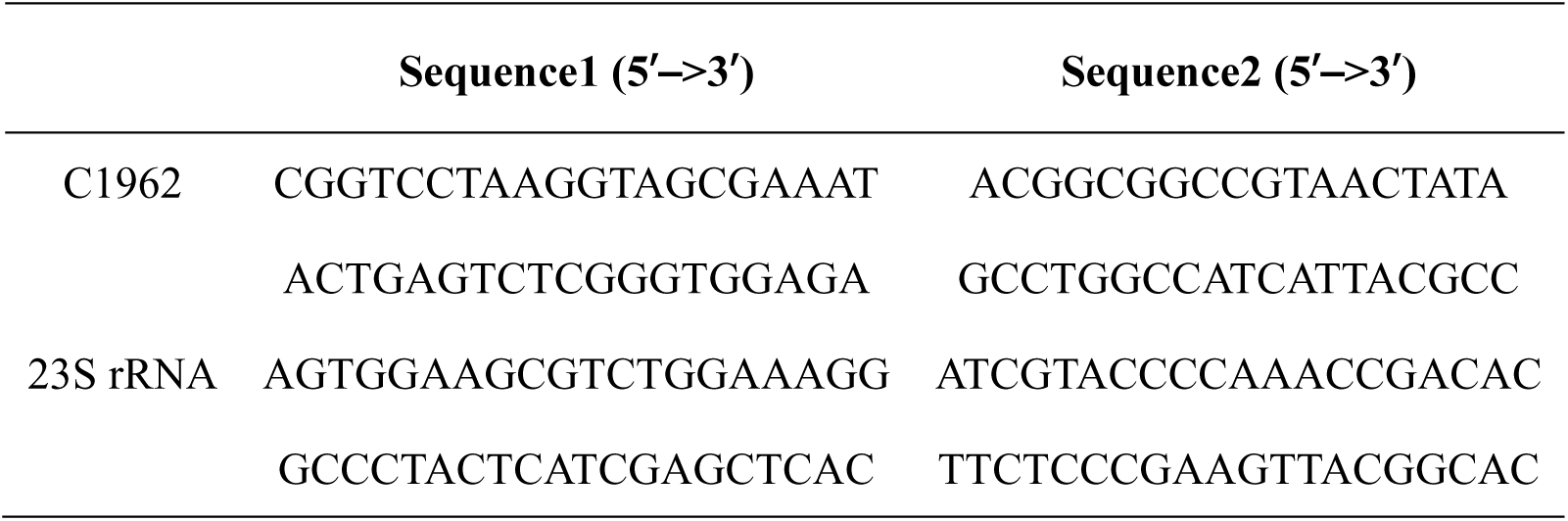

### Isolation and antimicrobial susceptibility test of E. coli

The soil samples were dissolved in phosphate-buffered saline, and large particles were removed using static precipitation. The supernatant was plated into MacConkey agar, followed by incubation at 37°C for 18 h. The pink, round, medium-sized colonies were picked as *E. coli* colonies. The antimicrobial susceptibility of the *E. coli* strains was determined by the Kirby-Bauer assay. Briefly, *E. coli* was inoculated into Mueller-Hinton agar (Sigma Aldrich, USA), and 30 µg kanamycin K (Liofilchem, Italy) was used for the antimicrobial susceptibility test. When the zone diameter was less than 13 mm, *E. coli* was considered resistant to kanamycin. The IMViC test kit (HiMedia, India) was employed to confirm the *E. coli* strains.

Urine samples were obtained from 8 healthy volunteers. 20 mL urine was centrifuged by 8000 g for 10 minutes and the precipitate was resuspended in 1 mL PBS. The method of isolation and antimicrobial susceptibility test consistent with the above. The study was approved by the Ethical Committee of Fujian Medical University, Fuzhou, China (2022-120).

The minimal inhibit concentrations (MICs) of WT and RlmI^K201A^ mutants were determined by ETEST (kanamycin 0.016-256 µg/mL; Liofilchem).

### Measuring the relative mRNA levels of methyltransferase

*E. coli* strains such as WT, *dgcZ*^+^ and *dgcZ* ^G206A,G207A^ were cultivated in LB medium at 37°C overnight and transferred to Vogel-Bonner medium at a 1:1000 ratio with 10 mM acetate at 25°C for 12 h and then induced with 0.2 mM IPTG at 25°C for 20 h. Ten OD cells were harvested for RNA extraction using an RNA extraction kit (Sangon Biotech, China). Real-time RT-PCR was performed using a reverse transcription kit and real-time PCR kit (Applied Biosystems, USA) with three replicates. The following primers were used.

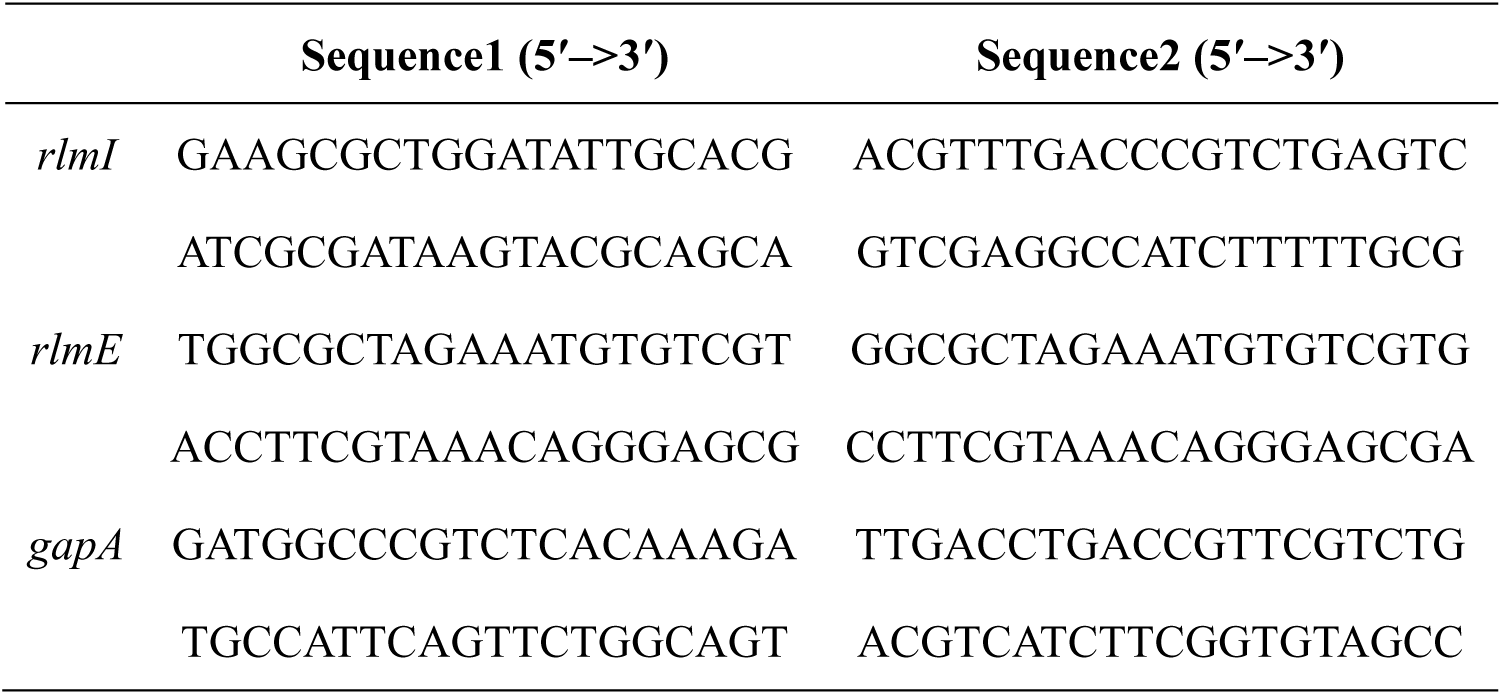

### Analysis of ribosomal subunits by sucrose density gradient centrifugation

*E. coli* strains were cultivated in LB medium at 37 °C overnight and transferred to Vogel-Bonner medium at a 1:1000 ratio with 10 mM acetate and 3 μg/mL kanamycin for 25°C for 24 h. Twenty OD cells were harvested and repeatedly freeze‒thawed three times. The cells were treated with 1 mL of lysis buffer (20 mM HEPES, 0.5 mM MgCl_2_, 200 mM NH_4_Cl, 1 mg/mL lysozyme, 50 units/mL benzonase, pH 7.5) at 4°C for 20 min with vigorous shaking. After lysis and centrifugation at 10,000 × *g* for 10 min at 4°C. The supernatant lysate was layered on top of a sucrose gradient (10%-50%, *w*/*v*) in lysis buffer and separated by ultracentrifugation in a Beckman SW-28 Rotor at 120,000 × *g* for 12 h at 4°C. The suspension was recovered to 25 components and quantified by measuring the absorbance at 260 nm on a NanoDrop 2000 spectrophotometer.

### Molecular dynamics simulations

The computational tool has been microsecond-scale molecular dynamics (MD) simulation ^24^. In MD simulation, the corresponding Newton’s equations of motion are integrated numerically after choosing reliable force fields, allowing us to monitor each individual atom in the system in a wide variety of setups, including liquids at interfaces, in solid walls or biological membranes, among others^25,26^. The number of particles and the pressure and temperature of the system were fixed, whereas the volume was adjusted accordingly. MD simulation can model hydrogens at the classical or quantum levels^27^. Besides its energetic and structural properties, it provides access to time-dependent quantities such as diffusion coefficients or spectral densities, enhancing its applicability. In the present study, we conducted MD simulations of c-di-GMP bound at rRNA methyltransferases (RlmI). The simulation system contained one rRNA methyltransferase and 7 c-di-GMP molecules fully solvated by 21,818 TIP3P water molecules in potassium chloride solution at the 0.15 M concentration, yielding a system size of 68,532 atoms. In each of the two statistically independent MD simulations, one single c-di-GMP molecule was placed at the center of the simulation box, near domains I, II and III of RlmI, whereas the remaining 6 free c-di-GMPs were randomly distributed around the RlmI protein according to the default settings of Charmm-GUI “Multicomponent Assembler”. All MD simulation inputs were generated using the CHARMM-GUI platform ^28,29^, and the CHARMM36m force field ^30^ was adopted for interactions between rRNA methyltransferases and c-di-GMP. The force field used also included the parameterization of the species c-di-GMP. All bonds involving hydrogens were set to fixed lengths, allowing fluctuations of bond distances and angles for the remaining atoms. The crystal structure of rRNA methyltransferases was downloaded from RCSB PDB Protein Data Bank (file name ”3c0k”). The system was energy-minimized for 50,000 steps and well-equilibrated (NVT equilibration in Figure 16 of SI) for 250 ps before the production of MD simulation. Production runs were performed with an NPT ensemble for 2 µs. The pressure and temperature were set at 1 atm and 310.15 K, respectively, to simulate the human body environment. The GROMACS 2021 package was employed in all MD simulations ^31^. Time steps of 2 fs were used in production simulations, and the particle mesh Ewald method with a Coulomb radius of 1.2 nm was employed to compute long-range electrostatic interactions. The cutoff for Lennard‒Jones interactions was set to 1.2 nm. The pressure was controlled with a Parrinello-Rahman piston with a damping coefficient of 5 ps^−1^, whereas the temperature was controlled using a Nosé-Hoover thermostat with a damping coefficient of 1 ps^−1^. Periodic boundary conditions in three directions of space were considered. We employed the “gmx-sham” tool of the GROMACS 2021 package to perform the Gibbs free energy landscape analysis. Moreover, the software VMD ^32^ and UCSF Chimera^33^ were used for trajectory analysis and visualization.

The radius of gyration *R*_g_, used as a reaction coordinate in the computation of Gibbs free energy landscapes, was determined as follows:

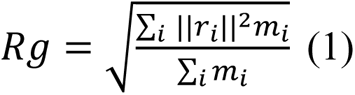

where *m*_i_ is the mass of atom *i*, and *r*_i_ is the position of the same atom with respect to the center of mass of the selected group. RMSD was calculated as follows:

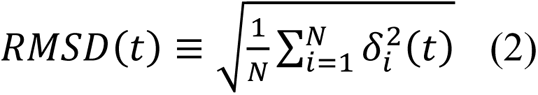

where *δ*_i_ is the difference in distance between atom *i* [located at *x*_i_(*t*)] of the catalytic domain and the equivalent location in the crystal structure.

RMSF values were obtained as follows:

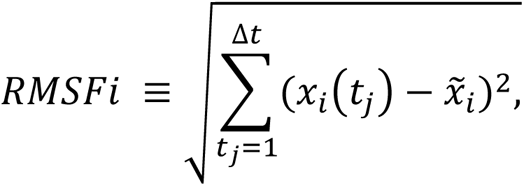

where 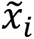 is the time average of *x*_i_, and Δ*t* is the time interval at which the average was taken.

### ITC assay

In the ITC assay, c-di-GMP (#C057 of Biolog), RlmI, and its mutants were prepared in titration buffer (20 mM Tris, 50 mM NaCl, 200 mM KCl, pH 7.0). Protein concentrations were measured based on Coomassie brilliant blue staining. The ITC titrations were performed using a MicroCal iTC200 system (GE Healthcare, PA, USA) at 25°C. Each titration process involved 22 injections with 5 μL c-di-GMP. The stock solution of c-di-GMP at 0.5 mM was titrated into WT or mutant RlmI (25 μM) in sample cells of 200 μL volume individually. c-di-GMP was titrated into 200 μL titration buffer as a control for subsequent data processing. The resulting titration curves were processed using the Origin 7.0 software program (OriginLab) according to the “one set of sites” fitting model.

### Streptavidin blotting assay

In this assay, RlmI (0.1 mg/mL) and its mutants were incubated with 10 μM biotin-c-di-GMP in a reaction buffer (20 mM Tris, 50 mM NaCl, 200 mM KCl, pH 7.0) at 37°C for 1 h. These samples underwent UV-crosslinking on ice for 0.5 h to further link c-di-GMP to RlmI. These linked samples were divided into two parts for analysis by western blotting. After incubation with IRDye 800CW Conjugated Streptavidin (#926-32230; LI-COR Biosciences, USA) at room temperature for 2 h, another membrane was incubated with anti-His antibody (05-949, Millipore, USA) at 4°C for 12 h and then incubated with an IRDye 800 secondary antibody for 1 h. The resulting membranes were visualized with an Odyssey Infrared Imaging System (LI-COR Biosciences).

### Isolation and quantification of c-di-GMP in E. coli

The c-di-GMP isolation was performed as discussed previously^6,34^. Briefly, *E. coli* cells with 50 OD were harvested and resuspended in 2 mL of ddH_2_O. Further, 8 mL of the mixture containing 50% methanol and 50% acetonitrile was added to extract intracellular c-di-GMP. Meanwhile, 1 μM cGMP was added as the internal reference. For c-di-GMP absolute quantification, the density of *E. coli* sediment is defined as 1 mg/mL. The extracts were analyzed by ultra-high-performance liquid chromatography coupled with ion mobility mass spectrometry (UPLC-IM-MS). UPLC-IM-MS was performed using a Waters UPLC I-class system equipped with a binary solvent delivery manager and a sample manager coupled with a Waters VION IMS Q-TOF mass spectrometer equipped with an electrospray interface (Waters Corporation, CT, USA).

### Determination of the strain growth curve in Vogel-Bonner medium

As discussed above, the strains WT, Δ*dgcZ*, *rlmI*^K201A^ and Δ*dgcZ rlmI*^K201A^ were grown in Vogel-Bonner medium with 10 mM acetate at 25°C. For kanamycin treatment, 0, 1.5, 3, 6 and 9 μg/mL kanamycin was added to the Vogel-Bonner medium, respectively. These cell concentrations were measured at OD_600_ using NanoDrop 2000 spectrophotometer at 8, 12, 16, 24, and 32 h. Then, the growth curve was drawn using GraphPad Prism 6.

### Statistical analysis

Pairwise comparisons were performed using two-tailed Student’s *t*-test, and statistical significance was set at \**p* < 0.05, \*\**p* < 0.01. Error bars represent the mean ± range.

## Results

### Ribosomal RNA large subunit methyltransferases are c-di-GMP effectors

In a previous study, we screened c-di-GMP-binding proteins in *E. coli* using a proteomic microarray and found that the rRNA

methyltransferases RlmI and RlmE were potential c-di-GMP effectors^6^ **(Fig. 1A)**. Based on the demonstration of binding between c-di-GMP and the methyltransferases, we speculated that c-di-GMP might affect rRNA methylation activity. To test this hypothesis, we examined the activity of four methyltransferases in the presence of c-di-GMP, using rRNA methylation as our indicator. For this purpose, we synthesized unmethylated 23S rRNA positions 1932-1991 and 2522-2581 for m5C1962 by RlmI ^35^ and m2U2552 by RlmE ^36^, respectively. The methyltransferases can catalyze the production of methylated rRNA and exhibit specific peaks in HPLC **(Fig. 1B-C)**. However, when c-di-GMP was introduced, methylation activities were significantly inhibited in a dose-dependent manner. We compared the effect of additional c-di-GMP on methylation products, using the group without c-di-GMP treatment as the reference value. Specifically, 5 μM c-di-GMP inhibited the activity of RlmI and RlmE by 49% and 31%, respectively **(Fig. 1D)**.

**Figure 1.**
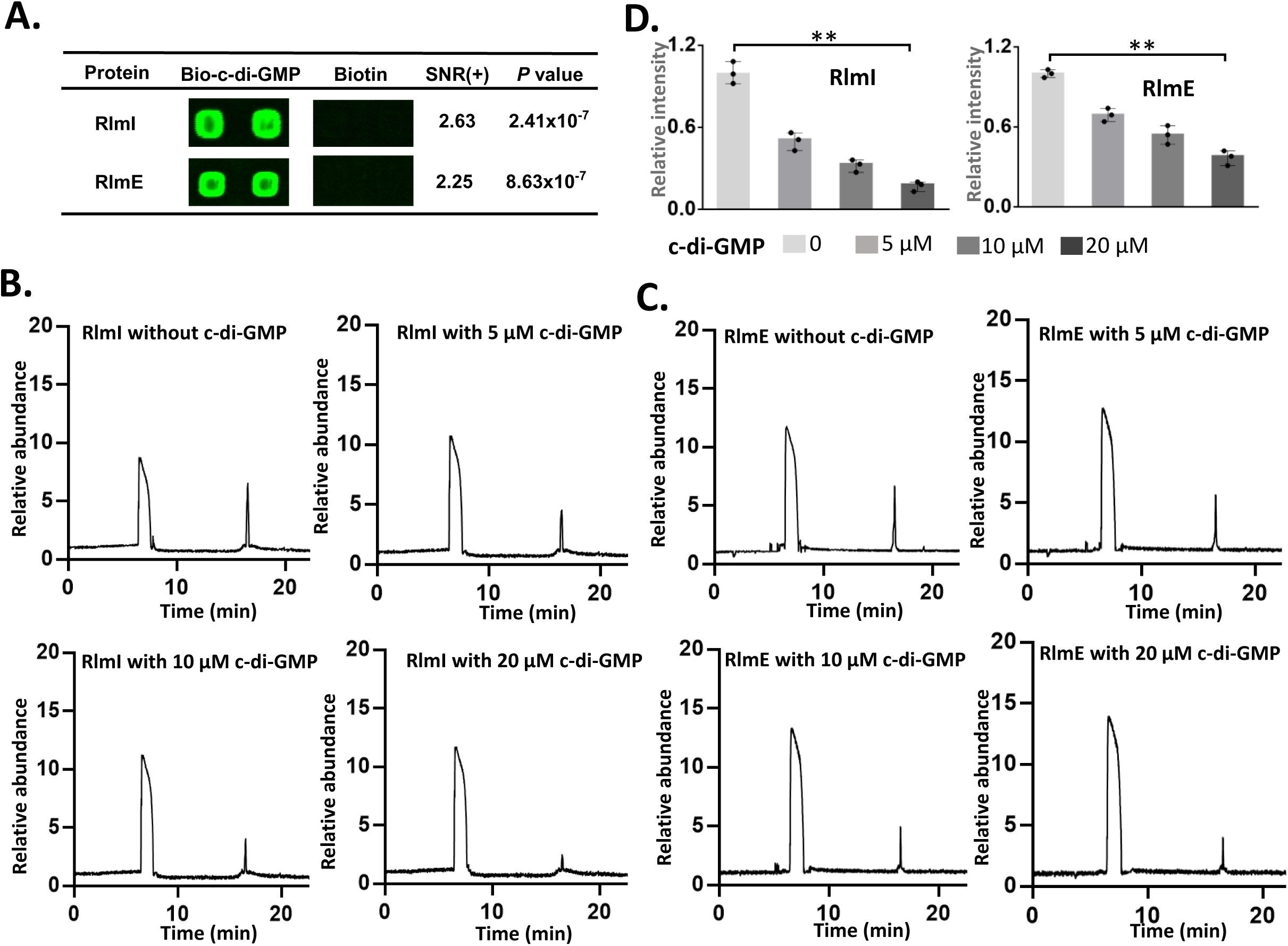
Ribosomal RNA large subunit methyltransferases are c-di-GMP effectors. (A) *E. coli* proteome microarrays were probed with biotin-c-di-GMP and biotin. Obvious binding differences of RlmI and RlmE on the microarrays incubated with biotin-c-di-GMP and biotin were observed. Two spots per protein were observed, and the positive signal-to-noise ratio [(SNR) (+)] represented the average SNR of the two duplicate spots. (B-C) *In vitro* methylation reaction. The synthesized rRNA fragments were used for *in vitro* methylation enzyme activity testing. The HPLC peaks are derived from RlmI (B) and RlmE (C) with the treatment of 0, 5, 10 and 20 μM c-di-GMP, respectively. (D) Quantitative results of HPLC peak. The methylated rRNAs were detected by HPLC and quantified by area under the curve. The bar chart shows the relative enzyme activity with the data points, using the reaction without the addition of c-di-GMP as the baseline (three preparations, mean ± range; \*\**p* < 0.01, two-tailed Student’s t-test).

### c-di-GMP inhibited ribosome assembly, with RlmI as the main effector

rRNA methylation is a prerequisite for the accurate assembly of ribosomes. We hypothesized that c-di-GMP might affect ribosomal assembly in *E. coli* (*E. coli* BW25113 as reference strain) by inhibiting methylation activity. To investigate the regulatory role of c-di-GMP, we constructed the strains of *dgcZ* knockout and overexpression. Compared with wild-type (WT) strains, the c-di-GMP level in *dgcZ* overexpressing strains increased by 12.2 times **(Fig. S1A)**. Then, we used a sucrose density gradient (SDG) assay to detect the ribosome particle, and the results showed that both knockout and overexpression of *dgcZ* could not affect the ribosome abundance of 50S compared with WT strain without antibiotic treatment **(Fig. 2A)**. c-di-GMP is a stress response factor. Therefore, we hypothesized that the regulation of ribosome assembly by c-di-GMP might occur under antibiotic stress. We treated *E. coli* cells with kanamycin, a ribosome-targeted antibiotic that can also elevate the cellular c-di-GMP level in *E. coli* by increasing the mRNA levels of *dgcZ*^6,37^. This elevation was controlled by the RNA-binding protein *csrA* ^38,39^.

**Figure 2.**
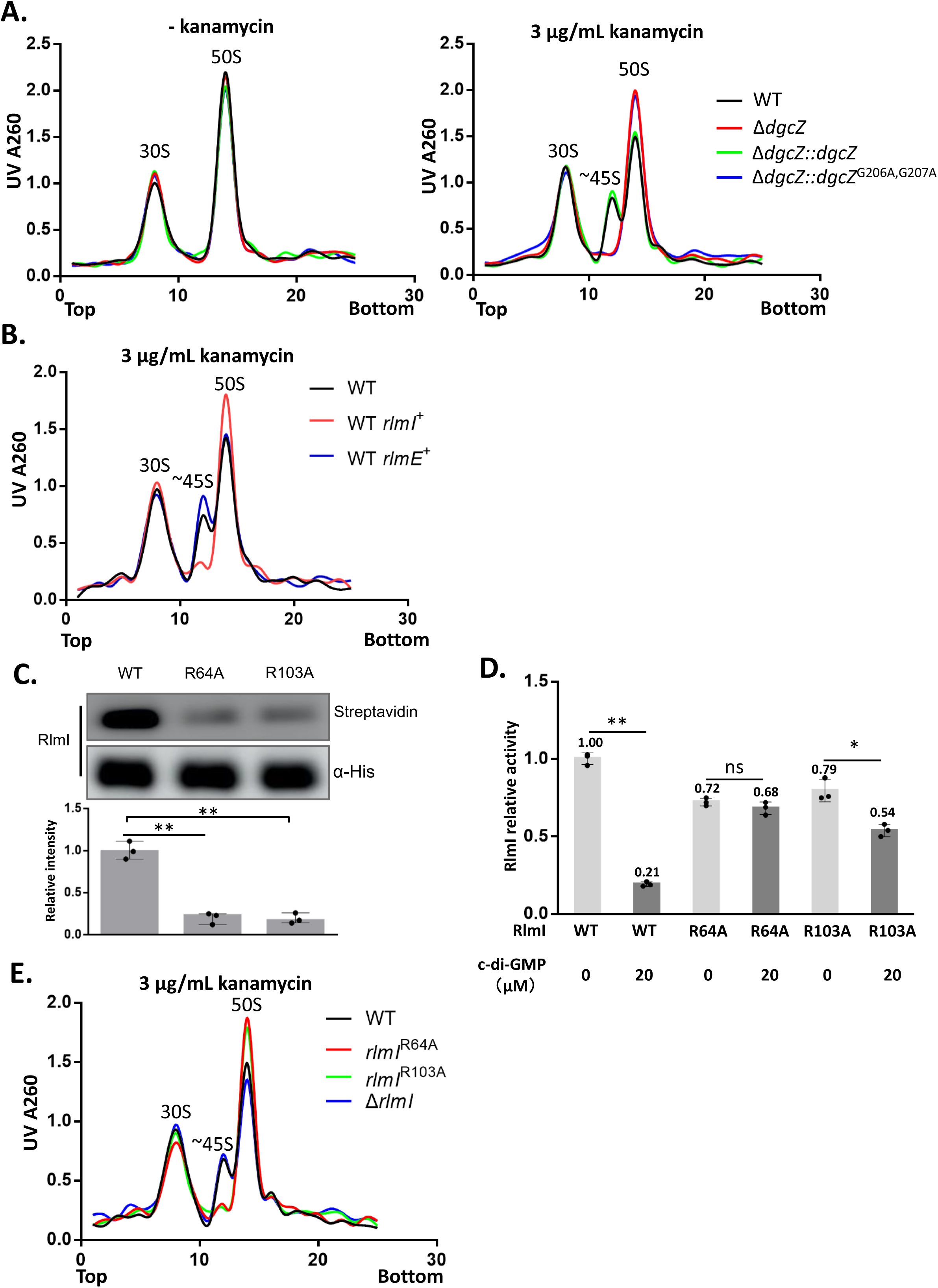
c-di-GMP inhibits ribosome assembly, with RlmI as the main effector. (A) SDG assay for the strains with elevated c-di-GMP. c-di-GMP was elevated by treatment with kanamycin or overexpression of DgcZ, and the ribosome particle was assayed by SDG. The corresponding three peaks represent the ribosome particles of 30S, pre-50S, and 50S. (B) SDG assay for the strains overexpressing two methyltransferases. RlmI and RlmE were overexpressed under the conditions of kanamycin treatment, and the ribosome particle was assayed by SDG. (C) Streptavidin blotting assays for WT and mutant RlmI. The arginine on RlmI was mutated to alanine, and the interaction of c-di-GMP and RlmI mutants was determined. The results indicated that R64A and R103A weakened the binding of c-di-GMP and RlmI. Streptavidin represents the interaction signals, and α-His represents the protein levels. The bar chart shows the relative intensity of streptavidin with the data points (three preparations, mean ± range; \*\**p* < 0.01, two-tailed Student’s t-test). (D) *In vitro* methylation assay of the two RlmI mutants. The synthesized rRNA fragments were used as substrates, and the reaction products were analyzed by HPLC. The bar chart shows the relative activity of RlmI with the data points (three preparations, mean ± range; ns: no significant difference, \**p* < 0.05, \*\**p* < 0.01, two-tailed Student’s t-test). (E) SDG assay for the strains with RlmI depletion and mutation. Strains with endogenous depletion and mutation of RlmI were constructed using the Red-recombination system. These strains were treated with kanamycin, and the ribosome particle was assayed by SDG.

Following kanamycin treatment, the c-di-GMP concentrations in WT cells were 6.2-fold higher than in untreated cells **(Fig. S1A)**. Notably, the c-di-GMP levels in *dgcZ*-defective cells were not responsive to kanamycin **(Fig. S1A)** because DgcZ serves as an effector for kanamycin-induced elevation of c-di-GMP levels. An SDG assay revealed that the ribosome disintegrated into 30S and 50S particles at low Mg^2+^ concentrations, and ∼45S particles^11^ were observed in kanamycin-treated WT cells and *dgcZ* replenishment strain (Δ*dgcZ::dgcZ*) **(Fig. 2A)**. However, the strain with *dgcZ* inactivation mutation did not show the presence of 45S particles (Δ*dgcZ::dgcZ*^G206A,G207A^). The results indicated that the increase in c-di-GMP levels inhibited the assembly of ribosomal large subunits in *E. coli*. In addition, we found c-di-GMP inhibited the activity of four methyltransferases and downregulated the methylation of 23S RNA *in vitro* **(Fig. 1D)**. Then, the four methyltransferases were overexpressed in kanamycin-treated WT cells to examine the role of the methylation enzymes in c-di-GMP-regulated ribosome assembly. We found that the overexpression of the methylases did not affect c-di-GMP levels **(Fig. S1B),** and the overexpression of RlmI weakened the effect of c-di-GMP on ribosomal assembly **(Fig. 2B)**. Thus, RlmI was the major effector of c-di-GMP in regulating ribosome assembly.

c-di-GMP binds to its effectors via Arg residues^40^. We mutated all Arg residues to Ala in RlmI to determine the binding sites on RlmI. Then, we developed the *in vitro* assay in which purified RlmI mutants were incubated with biotin-c-di-GMP, ultraviolet (UV)-crosslinked, and probed with fluorescent streptavidin ^41,42^. We observed that RlmI mutants with R64A and R103A exhibited a significantly weakened interaction with c-di-GMP **(Figs. 2C and S2)**. Furthermore, when we determined the activity of RlmI mutants, both RlmI^R64A^ and RlmI^R103A^ displayed methylation activities slightly lower than that of RlmI under 5 μM rRNA substrate. Upon treatment with 20 μM c-di-GMP, the methylation activity decreased by 80%, 6%, and 32% in RlmI, RlmI^R64A^, and RlmI^R103A^, respectively **(Fig. 2D)**. The effect of c-di-GMP for RlmI was significantly weakened under R64A and R103A mutations in comparison to that of WT, which validated that R64 and R103 are the key sites to the binding of c-di-GMP to RlmI.

We eliminated the effect of c-di-GMP on RlmI by mutation of the binding sites R64A and R103A to validate whether RlmI was the main effector of c-di-GMP in ribosome assembly. With kanamycin treatment, WT, *rlmI^R64A^*, *rlmI^R103A^,* and △*rlmI* cells were assayed by SDG. Under kanamycin stimulation, elevated c-di-GMP in WT cells inhibited RlmI, resulting in missing rRNA methylation. This inhibition of RlmI impairs ribosomal assembly and leads to the production of 45S precursors. The *rlmI* knockout strains serve as a positive control for demonstrating *rlmI* functional defects. In addition, the R64A and R103A mutants abolished the effect of c-di-GMP on RlmI. The results showed that the ∼45S particle was observed in WT and Δ*rlmI* cells but not in R64A and R103A mutants **(Fig. 2E)**. Thus, RlmI played a key role in ribosomal assembly under kanamycin stress, and c-di-GMP regulated its activity.

### c-di-GMP induces the closure of the catalytic pocket of RlmI

To reveal the structural mechanism of c-di-GMP regulating RlmI enzyme activity, we investigated the conformational changes of RlmI during its interaction with c-di-GMP in an aqueous ionic solution using molecular dynamics (MD) simulation. In MD simulation, root mean square deviation (RMSD) indicated the fluctuations and stability of the conformations of RlmI and root mean square fluctuation (RMSF) revealed flexibility during the full simulation period. By RMSD and RMSF, we found that residues 160-170, 302-320, and 370-390 were mainly involved in the conformational fluctuation of RlmI **(Fig. 3A)**. We labeled residues 160-170 as “Domain-I” (DM-I), residues 302-320 as “Domain-II” (DM-II), and residues 370-390 as “Domain-III” (DM-III). DM-I and DM-III are the regions of the protein that correspond to the RNA-binding area, whereas DM-II is located near the S-adenosyl-L-methionine (AdoMet) binding-related area. An overall view of the evolution of RlmI fluctuations revealed a distinct conformational fluctuation around 0.8 µs during the simulation (average of Trajectory #1 and Trajectory #2) **(Fig. 3B)**. Combining RMSD, RMSF, and trajectory analysis results, the RlmI was found to exist in two states, State-I and State-II, while interacting with c-di-GMP. c-di-GMP can interact with the RNA-binding area (State-I), and the domain DM-III shut down after 0.8 µs. The results suggested that (1) c-di-GMP can interact with the RNA-binding domains and then induce the closure of DM-III, and (2) it was corroborated that the “on-off” of the RNA-binding area was mainly embodied by DM-III to a large extent **(Fig. 3C and Video 1-2)**. The dynamic process of c-di-GMP-induced conformational rearrangements in the active domain of RlmI was similar to that of other c-di-GMP effectors such as YcgR ^43^, FleQ ^44^, and CheR1 ^45^.

**Figure 3.**
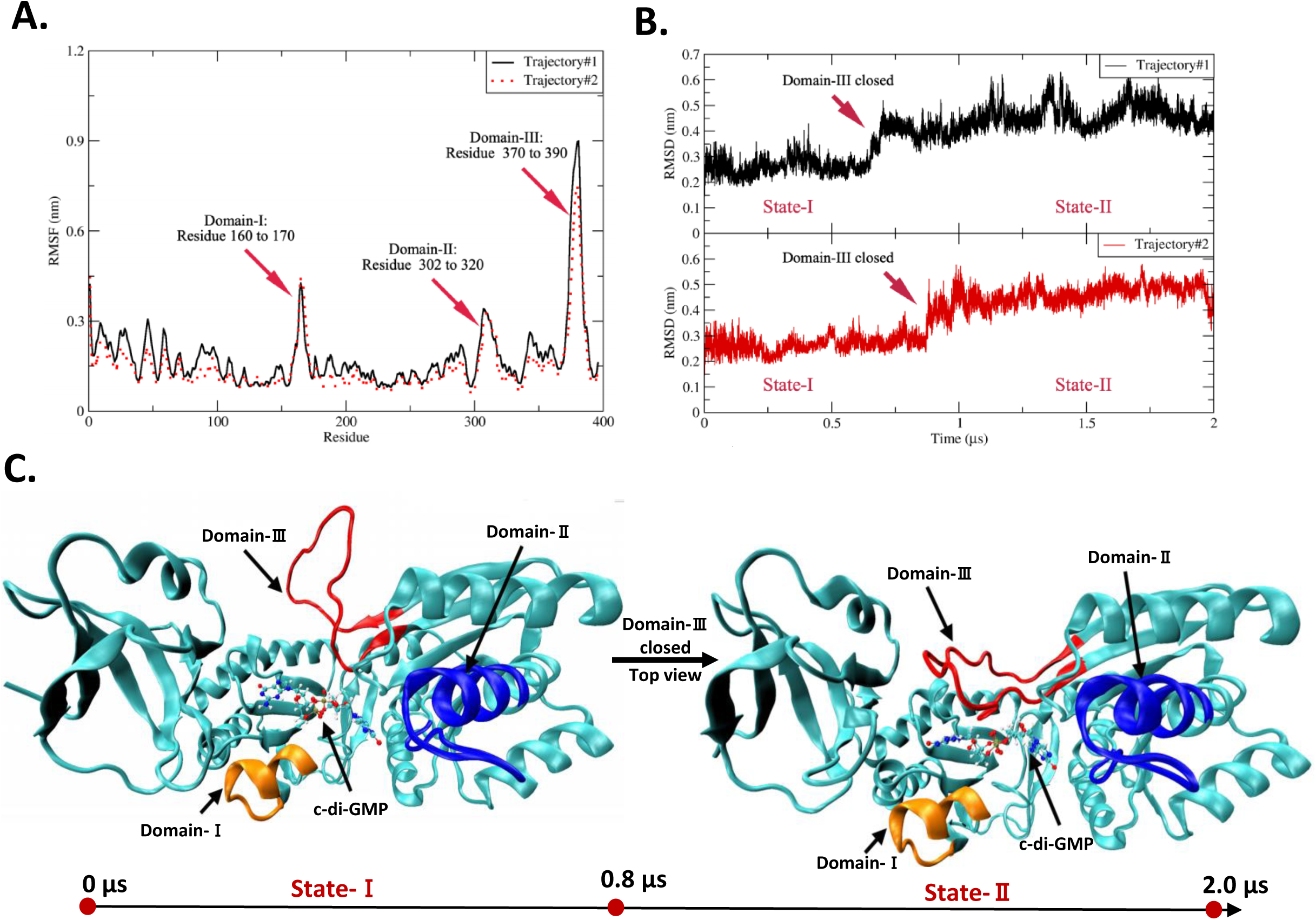
Binding model of c-di-GMP and RlmI. (A) RMSD value of RlmI during interaction with c-di-GMPs. The arrow marks the main changes at the residue level. The solid and dashed lines represent two simulated trajectories, and the degree of overlap represents the convergence of two trajectories. (B) RMSF value of RlmI during interaction with c-di-GMP. The arrow marks the main changes on the timeline. (C) Representative snapshots of the transition between RlmI State-I and State-II. The arrow marks the main changes in DM-III.

### R64, R103, G114, and K201 residues of RlmI interact with bound c-di-GMP

We employed Gibbs free energy analysis to identify the dominant conformation of RlmI and c-di-GMP complex. The Gibbs free energy surfaces were shown for the two runs and their average **(Fig. 4A),** using RMSD and radius of gyration as the variables. We identified the free energy basin, the one with the lowest free energy (set to 0 kJ/mol) **(Fig. 4A, yellow point)**, and found that the corresponding regions were almost overlapping for the three sets (**Fig. 4B)**. Thus, the results indicated the two independent simulated trajectories as convergent and physically equivalent.

**Figure 4.**
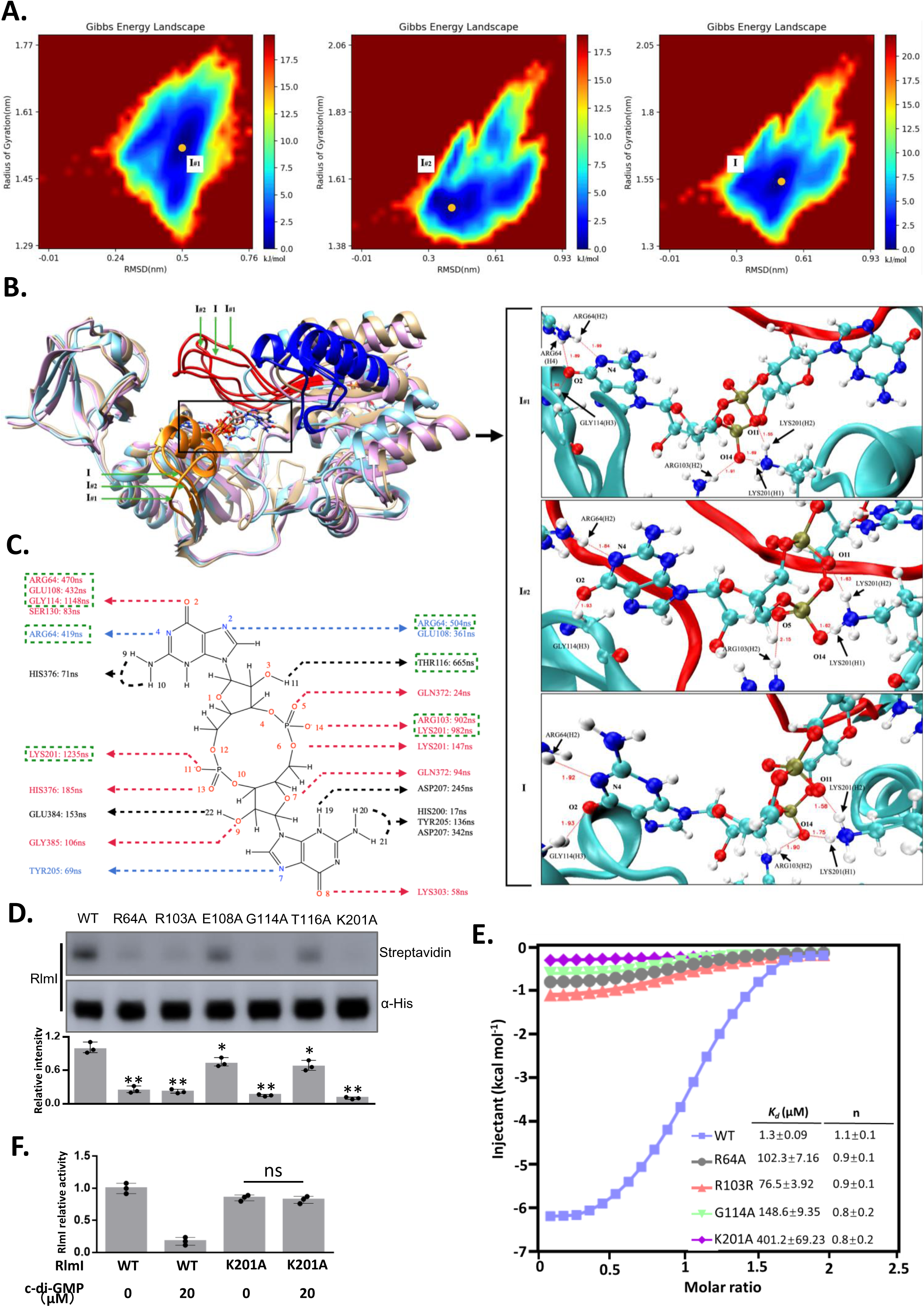
Determination of the binding sites of c-di-GMP on RlmI. (A) Gibbs free energy landscapes illustrating the binding of c-di-GMP to RlmI. I#1, I#2, and I represent trajectories 1, 2, and average, respectively. (B) Interaction model of c-di-GMP and RlmI. The key binding sites R64, R103, G114, and K201 are highlighted. I#1, I#2, and I represent trajectories 1, 2, and average, respectively. (C) Interaction landscapes of c-di-GMP and RlmI. The interaction sites between c-di-GMP and RlmI in MD simulation are marked in the plane diagram of c-di-GMP. The annotated duration represents the interaction time between c-di-GMP and RlmI in the 10 μs MD simulation, with green boxes highlighting residues with interaction times greater than 400 ns. (D) Streptavidin blotting assays comparing WT and mutant RlmI. The potential binding sites within RlmI were mutated to alanine, and the interaction with c-di-GMP was determined. Streptavidin represents the interaction signals, and α-His represents the protein levels. The intensity of WT was used as the reference for statistical analysis of the variants. The bar chart shows the relative intensity of streptavidin with the data points (three preparations, mean ± range; \**p* < 0.05, \*\**p* < 0.01, two-tailed Student’s t-test). (E) ITC analysis of the binding between c-di-GMP and RlmI. c-di-GMP was titrated into RlmI and its mutants by three repetitions. (F) *In vitro* methylation assay of the RlmI mutants. The bar chart shows the relative activity of RlmI with the data points (three preparations, mean ± range; ns: no significant difference, two-tailed Student’s t-test).

To explore the binding sites of c-di-GMP and RlmI, we superimposed the stable-state configurations of RlmI and c-di-GMP for the three sets **(Fig. 4B)**. Two independent trajectories #1 and #2 were taken into consideration for the computational analysis and the average was selected for convergence and physical equivalence analysis. The average conformation showed that R64 and G114 together stabilize the guanosine moiety of c-di-GMP. Correspondingly, R103 and K201 act to stabilize the negatively charged region of c-di-GMP. It is evident that R103 and K201 formed a stable hydrogen bond with the oxygen atom of the phosphate group of c-di-GMP.

Noncovalent interactions, including hydrogen bonds, coordination bonds, and salt bridges, are crucial for maintaining the tertiary structure of proteins. Benefiting from the all-atom-level precision of molecular dynamics simulations, we analyzed the hydrogen bond interaction map of c-di-GMP with RlmI using time-dependent atomic site distances between selected atomic sites to uncover the interaction mode of c-di-GMP with RlmI, providing guidance for further experimental verification. Atomic detail sketches of c-di-GMP and the main residues described in this section are provided in **Figures S3 and S4**. While labeling the amino acid residues in the hydrogen bond interaction map of c-di-GMP with RlmI, we also labeled the lifetime of hydrogen-bonding interactions between c-di-GMP and corresponding amino acid residues. Considering that our molecular dynamics simulation spanned a timeframe of 2 µs, we subsequently performed site mutation verification on residues with hydrogen bond interaction lifetimes exceeding 400 ns. Six amino acid residues from RlmI were selected as the potential binding sites for c-di-GMP: R64, R103, E108, G114, T116, and K201 **(Fig. 4C)**.

Atom-atom distances as a function of time and bond lifetimes are presented in **Fig. S5-S15**.

We employed the streptavidin blotting assays to determine the interaction between c-di-GMP and RlmI mutants, aiming to validate the results obtained from the molecular dynamics simulations. The result revealed that amino acid residues R64, R103, G114, and K201 were crucial for the binding of c-di-GMP to the RlmI. In addition, E108A and T116A of RlmI slightly affected c-di-GMP binding **(Fig. 4D)**. We next performed isothermal titration calorimetry (ITC) titrations with these mutants and determined *K_d_* values of 1.3, 102.3, 76.5, 148.6, and 401.2 μM for RlmI, RlmI^R64A^, RlmI^R103A^, RlmI^G114A^, and RlmI^K201A^, respectively **(Figs. 4E and S16)**. We found that the stoichiometries have not significant different between RlmI and its variants, which showed 1:1 binding ratio with c-di-GMP **(Fig. 4E)**. Further, we observed that the activity of RlmI^K201A^ showed no significant change with 20 μM c-di-GMP treatment compared with c-di-GMP free group. **(Fig. 4F)**. The results suggested that R64, R103, G114, and K201 residues of RlmI were the critical sites for c-di-GMP binding.

### c-di-GMP regulates RlmI to promote antibiotic resistance

As a close correlation exists among c-di-GMP, ribosomal assembly, and antibiotic resistance^46,47^, we hypothesized that c-di-GMP regulated ribosomal assembly to promote antibiotic resistance by inhibiting RlmI activity. To test this hypothesis, we interfered with the interaction of c-di-GMP and RlmI by introducing the K201 mutation in endogenous RlmI (*rlmI*^K201A^), *dgcZ* depletion (△*dgcZ*), *dgcZ* overexpression (△*dgcZ dgcZ*^+^) and both mutants (△*dgcZ rlmI*^K201A^). Then, we determined the c-di-GMP level and methylation level of 23S rRNA C1962. The results showed that the cellular c-di-GMP levels and C1962 methylation levels did not significantly change in WT, △*dgcZ*, *rlmI*^K201A^, and △*dgcZ rlmI*^K201A^ without kanamycin treatment **(Figs. 5A** and **5B)**. However, the c-di-GMP levels of WT and *rlmI*^K201A^ increased about six times compared with △*dgcZ* and △*dgcZ rlmI*^K201A^ when treated by kanamycin, and the c-di-GMP levels of △ *dgcZ dgcZ*^+^ increased about eleven times compared with WT **(Fig. 5A)**. Furthermore, the methylation analysis showed that elevated c-di-GMP (WT, △ *dgcZ dgcZ*^+^) significantly decreased C1962 methylation, and the K210A mutation (*rlmI*^K201A^) abolished the effect of c-di-GMP **(Fig. 5B)**.

**Figure 5.**
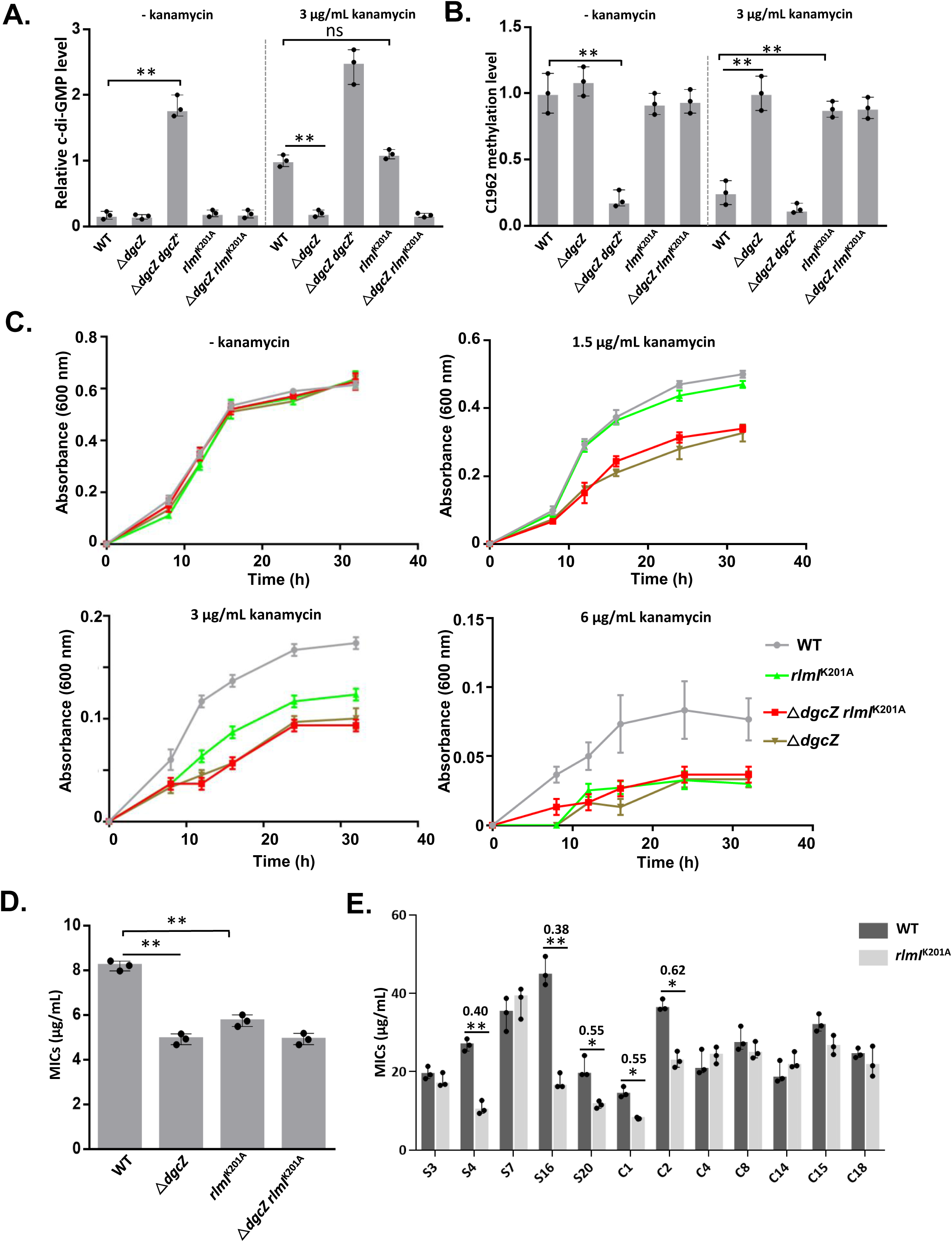
c-di-GMP regulates ribosome assembly to promote drug resistance. (A) Relative c-di-GMP level of WT, Δ*dgcZ*, Δ*dgcZ dgcZ*^+^, *rlmI*^K201A^ and Δ*dgcZ rlmI*^K201A^ strains. The intracellular c-di-GMP concentrations were determined by UPLC-IM-MS. The bar chart shows the relative quantification of c-di-GMP with the data points (three preparations, mean ± range; \*\**p* < 0.01, two-tailed Student’s t-test). (B) Methylation level of C1962 in 23S rRNA of WT, Δ*dgcZ*, Δ*dgcZ dgcZ*^+^, *rlmI*^K201A^ and Δ*dgcZ rlmI*^K201A^ strains. The endogenous methylation level in C1962 was determined by meRIP-qPCR in triplicate. The bar chart shows the relative quantification of methylated rRNA with the data points. (C) Growth curves of the WT, Δ*dgcZ*, *rlmI*^K201A^ and Δ*dgcZ rlmI*^K201A^ strains. The *E. coli* strains were cultured in Vogel-Bonner medium with 0, 1.5, 3 and 6 μg/mL kanamycin treatment, resectively. The growth was measured after 8, 12, 16, 20, 24, and 32 h in triplicate (three preparations, mean ± range). (D) The MICs for the WT, Δ*dgcZ*, *rlmI*^K201A^ and Δ*dgcZ rlmI*^K201A^ strains. The bar chart shows the MICs, which were determined by ETEST. The bar chart shows the value of MICs with the data points (three preparations, mean ± range; ns: no significant difference, \*\**p* < 0.01, two-tailed Student’s t-test). (E) The MICs of the resistant strains. The 12 kanamycin-resistant strains were mutated from RlmI K210 to alanine, and the MICs were determined by ETEST. The fold changes of MIC (MIC_*rlmI*^K201A^/MIC_WT) were annotated for the strains with statistically different MICs upon the mutation of K201 to Alanine. The bar chart shows the value of MICs with the data points (three preparations, mean ± range; \**p* < 0.05, \*\**p* < 0.01, two-tailed Student’s t-test).

We measured the growth curves and MICs of the four strains, *i.e.*, WT, Δ*dgcZ*, *rlmI*^K201A^, and Δ*dgcZ rlmI*^K201A^. We tested serial concentrations (0, 1.5, 3, 6 and 9 μg/mL) of kanamycin for strain growth curve. The strains cannot growth in treatment with 9 μg/mL kanamycin, so we determined the growth curve for other four concentrations. The curves showed that growth of the four strains were consistent when kanamycin was not added **(Fig. 5C)**. However, the growth of the Δ*dgcZ* strains decreased compared with that of the WT strains with 1.5, 3 and 6 μg/mL kanamycin treatment, indicating that c-di-GMP promoted the antibiotic tolerance of *E. coli*. Furthermore, we mutated the c-di-GMP binding site K201 of RlmI (*rlmI*^K201A^) and found that *rlmI*^K201A^ strains had greater sensitivity to antibiotics and slower growth rate compare to that of WT, which suggests that the interaction of c-di-GMP and RlmI enhances the antibiotic tolerance **(Fig. 5 C-D)**.

Kanamycin is an antibiotic that targets ribosomes and can endogenously increase the c-di-GMP level. Therefore, we speculate that c-di-GMP has a promoting effect on kanamycin resistance. To test this, we isolated 40 *E. coli* strains from soil (named S1-S20) and urine (named C1-C20) samples resistant to kanamycin and determined the cellular c-di-GMP levels to investigate the role of c-di-GMP and RlmI axis in kanamycin-resistant *E. coli*. The results revealed that 13/40 strains (S3, S4, S7, S12, S16, S20, C1, C2, C4, C8, C14, C15, C18) showed significant increase in c-di-GMP levels compared with the BW25113 strains **(Fig. S17A)**. Furthermore, we mutated the endogenous RlmI in K201 to alanine and determined the minimal inhibitory concentration (MIC). We obtained 12 of 13 K201 mutant strains, excluding the C12 strain, which might be a multidrug-resistant strain. Drug susceptibility results showed that the MICs of the S4, S16, S20, C1 and C2 strains substantially decreased following the mutation at K201 of RlmI **(Fig. 5E)**. Thus, c-di-GMP regulated RlmI activity to promote antibiotic tolerance, and this pathway played a key role in a portion of antibiotic-resistant *E. coli*.

### Effect of c-di-GMP on RlmI is conserved in multiple pathogenic bacteria

c-di-GMP is a ubiquitous bacterial secondary messenger, and RlmI is highly conserved in bacteria. Thus, we hypothesized that the binding and inhibition of c-di-GMP with RlmI from *E. coli* was the same for the RlmI homologues in other bacteria. RlmI protein sequences from a series of highly diverse bacteria were aligned with testing this hypothesis, and we found that the c-di-GMP binding region was reasonably well conserved in these bacteria **(Fig. 6A)**. Then, we selected *Salmonella typhimurium*, *Klebsiella pneumoniae*, and *Vibrio cholerae* as the exemplary members of this conserved set. We found that RlmI*^S. typhimurium^*, RlmI*^K. pneumoniae^*, and RlmI*^V. cholerae^* could bind to c-di-GMP and that the binding could be abolished when the lysine in RlmI was mutated **(Fig. 6B)**. Furthermore, the *in vitro* activity analysis showed that similar to RlmI*^E.coli^*, the aforementioned three RlmI homologues exhibited methylase activity for 23S rRNA and this activity could be inhibited by c-di-GMP **(Fig. 6C)**. Thus, the effect of c-di-GMP on RlmI is conserved in multiple pathogenic bacteria and might also play a role in antibiotic resistance.

**Figure 6.**
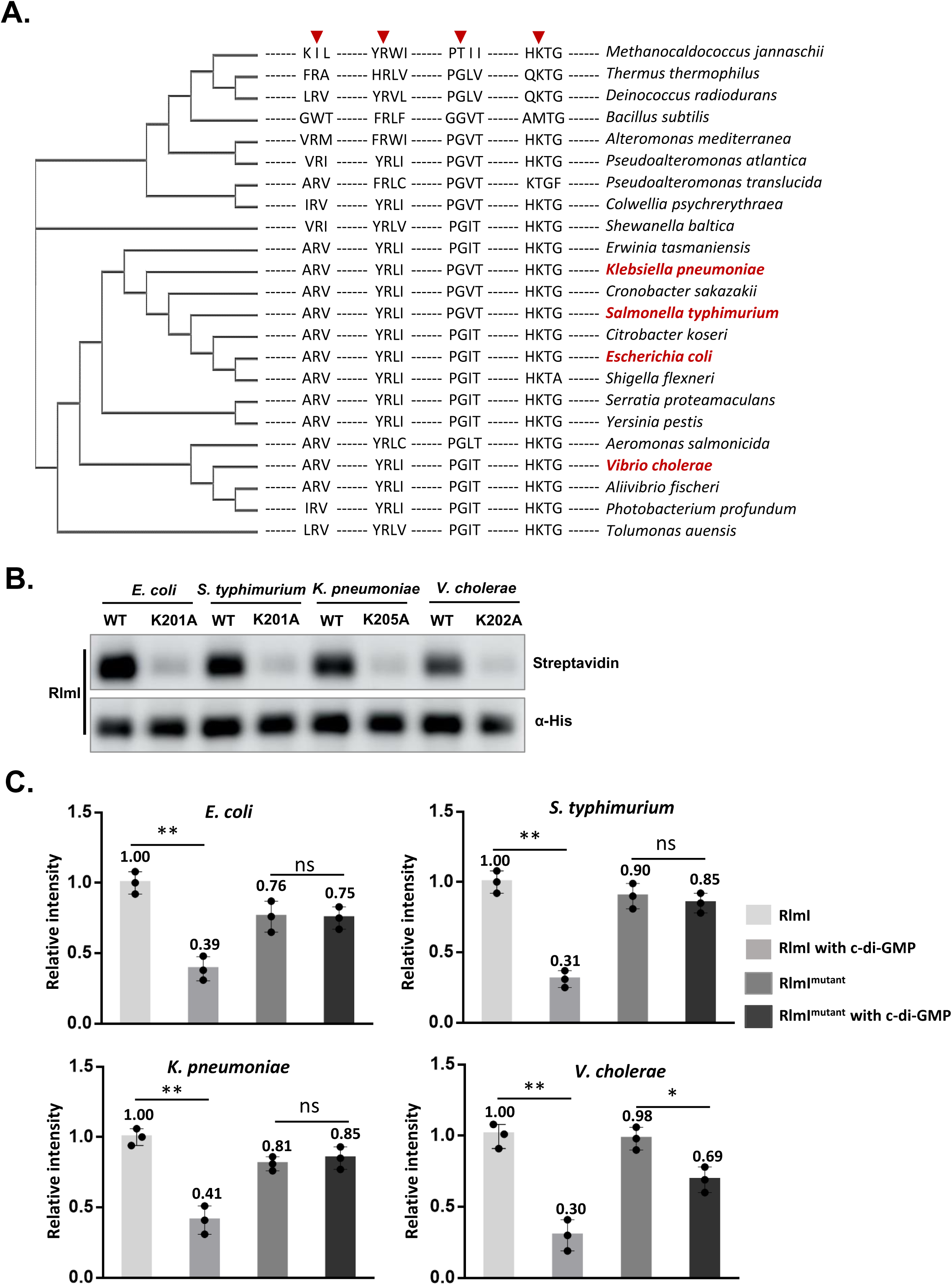
c-di-GMP binds RlmI and inhibits its activity conserved in multiple pathogenic bacteria. (A) CLUSTALW alignment of the binding sites of c-di-GMP and RlmI. Residues involved in c-di-GMP binding (R64, R103, G114, and K201) are marked by red arrows. (B) Streptavidin blotting assays for RlmI of four species. The WT and mutant RlmI interacted with biotin-c-di-GMP and were crosslinked by UV. Streptavidin and α-His present interaction signals and protein levels, respectively. (C) *In vitro* methylation assay for the RlmI of four species. The experiment was performed in triplicate. The bar chart shows the quantitative results of methylation products with the data points (three preparations, mean ± range; ns: no significant difference, \**p* < 0.05, \*\**p* < 0.01, two-tailed Student’s t-test).

## Discussion

c-di-GMP is a key secondary messenger in prokaryotes. rRNA methylation occurs in both prokaryotes and eukaryotes. This study discovered that c-di-GMP binds to four rRNA methyltransferases and inhibits their activities. RlmI was found to be the major effector of c-di-GMP in ribosome assembly. The molecular dynamics simulation analysis revealed the binding sites and models of c-di-GMP and RlmI. The MIC assay further demonstrated that c-di-GMP inhibits ribosome assembly, promoting antibiotic resistance in *E. coli*. Thus, we established a regulatory pathway among c-di-GMP and ribosome functions and revealed its role in antibiotic resistance.

Previous studies have reported that c-di-GMP regulates mature ribosome function through RimK in *Pseudomonas*^48,49^, EF-P in *Acinetobacter baumannii* ^50^, Vc2 riboswitches in *V. cholerae* ^51^. c-di-GMP regulates the glutamate ligase RimK, which catalyzes glutamate residues to the C-terminus of ribosomal protein RpsF to affect ribosomal function^48,49^. The binding of c-di-GMP enhances the function of EF-P, promoting translation efficiency and modulating bacterial physiology and virulence ^50^. Additionally, c-di-GMP binds to the Vc2 riboswitch, inducing structural changes that result in the switch-OFF and switch-ON states of translational initiation ^51^. This study revealed that c-di-GMP affects ribosome assembly, offering a new perspective on c-di-GMP’s role in ribosome regulation. Numerous accessory factors play a role in guiding the ribosome assembly process, including GTPases, rRNA modification enzymes, helicases, and maturation factors ^52^. Our findings establish c-di-GMP as an upstream regulatory signal for rRNA modification, creating a connection between environmental stimuli and ribosome function.

RlmI is a large ribosomal RNA subunit methyltransferase that methylates cytosine specifically at position 1962 (m5C1962) of 23S rRNA. In previous studies, RlmI depletion did not lead to abnormal ribosome assembly or growth arrest of *E. coli* at 20°C and 37°C^53^. Indeed, we found that RlmI depletion did not affect the ribosome abundance of 50S compared with WT strains when not treated with antibiotics.

However, with kanamycin treatment, the ∼45S particle was observed in △*rlmI* cells **(Fig. 2E)**. Thus, RlmI plays a key role in ribosomal assembly under kanamycin stress. As deletion of most ribosomal methyltransferases does not cause significant phenotypic changes, these studies have demonstrated that the function of methylases under different growth conditions may help understand the physiological significance of ribosome assembly.

rRNA methylation is a fundamental mechanism contributing to bacterial resistance against ribosome-targeting antibiotics. Genes encoding corresponding methyltransferases have been identified in clinical isolates of pathogenic bacteria ^54^. In 16S rRNA, methylation of C1405 and A1408 confer high-level resistance to aminoglycosides, which is catalyzed by the RsmF and Npm families, respectively. The phylogenetic analysis suggests that m^7^G1405 and m^1^A1408 methyltransferases share a common ancestor with aminoglycoside-producing soil bacteria. In 23S rRNA, the SAM methyltransferase family shares ancestry with the housekeeping RlmN methyltransferases, which incorporate the methylation of A2503 in 23S rRNA. However, the connection between RlmI and antibiotic resistance has not previously been reported. We also detected the transcriptional levels of RlmI for the drug-resistant strains and did not find any intrinsic change in expression levels **(Fig. S17B)**. Because RlmI is not the direct target of most of the known antibiotics, and its enzymatic activity could be regulated through the binding of c-di-GMP. We speculate that RlmI may function as an effector of the stress factor c-di-GMP, and the pathway of c-di-GMP regulating RlmI may play the role in intrinsic tolerance.

Bacteria mainly acquire resistance by altering resistance genes, reducing intracellular antibiotic concentrations, protecting antibiotic targets, and inactivating antibiotics^55^. c-di-GMP promotes the biofilm to reduce the permeability of antibiotics ^46,56^. Other mechanisms by which c-di-GMP promotes bacterial resistance remain to be studied. We found that the level of c-di-GMP increased in some drug-resistant *E. coli* and interfering with the effect of c-di-GMP on ribosome methylation in these

bacteria could increase their sensitivity to antibiotics. c-di-GMP has essential physiological functions in drug-resistant bacteria. However, not all drug-resistant bacteria have elevated intracellular c-di-GMP concentrations, and c-di-GMP regulates other drug-resistance-related pathways, such as biofilms. The question of whether the resistance pathway regulated by c-di-GMP has a synergistic or independent effect is worth exploring. Therefore, more data on drug-resistant strains are needed to examine the role of c-di-GMP in bacterial resistance.

In summary, we identified rRNA methyltransferases as the novel c-di-GMP effectors from *E. coli* proteome microarray assay. The functional analysis revealed an unexpected role of c-di-GMP in regulating ribosome assembly by inhibiting rRNA methylases, highlighting the physiological function of the regulatory axis in bacterial drug resistance.

## Acknowledgments

This study was supported by the National Natural Science Foundation of China (Grant No. 32000027), the Natural Science Foundation of Fujian Province, China (No. 2022J01197), Fourteenth Five-Year National Key Research and Development Program of China (2023YFC2307200), R&D Program of Guangzhou National Laboratory (No. GZNL2023A01005) and Natural Science Foundation of China (No. 92374110).

## Author contributions

Z.W.X., S.C.T. and J.M. conceived the idea. S.C.T. provided key reagent. S.Q.Y., X.R.L., J.Y.S., and H.C. performed protein mutation and interaction assay. X.T.X. performed the bacterial culture and protein purification. S.Q.Y., M.L., and X.T.X. performed an enzyme activity assay. X.T.X., X.R.L., X.T.X., and L.X.Z. performed the bacterial drug sensitivity test. Z.Y.H. and J.M. performed molecular dynamic simulation and structure analysis. Z.W.X., S.Q.Y., Z.Y.H., and J.M. prepared the figures. Z.W.X., S.Q.Y., Z.Y.H., and J.M. wrote the manuscript.

## Data and software availability

The crystal structure files, MD simulation files (input files, parameter files, topology files, etc.), and structures of c-di-GMP are available on the website https://github.com/Zheyao-Hu/RlmIcdiGMP. Moreover, all the software (free to use) packages used in this study were the official release version without any modification. The raw protein microarray data have been published in the Protein Microarray Database (www.proteinmicroarray.cn/) with the accession number PMDE226 (http://www.proteinmicroarray.cn/index.php/experiment/detail?experimen t_id=226).

## References

1 Ross, P., et al. Regulation of cellulose synthesis in Acetobacter xylinum by cyclic diguanylic acid. Nature 325, 279–281, doi:10.1038/325279a0 (1987).

2 Hengge, R. Principles of c-di-GMP signalling in bacteria. Nature reviews. Microbiology 7, 263–273, doi:10.1038/nrmicro2109 (2009).

3 Jenal, U., Reinders, A. & Lori, C. Cyclic di-GMP: second messenger extraordinaire. Nature reviews. Microbiology, doi:10.1038/nrmicro.2016.190 (2017).

4 Romling, U., Galperin, M. Y. & Gomelsky, M. Cyclic di-GMP: the First 25 Years of a Universal Bacterial Second Messenger. Microbiology and Molecular Biology Reviews 77, 1–52, doi:10.1128/mmbr.00043-12 (2013).

5 Obeng, N. et al. Bacterial c-di-GMP has a key role in establishing host–microbe symbiosis. Nature microbiology, doi:10.1038/s41564-023-01468-x (2023).

6 Xu, Z. et al. Interplay between the bacterial protein deacetylase CobB and the second messenger c-di-GMP. The EMBO journal 38, e100948, doi:10.15252/embj.2018100948 (2019).

7 Arai, T., Ishiguro, K., Kimura, S., Sakaguchi, Y. & Suzuki, T. Single methylation of 23S rRNA triggers late steps of 50S ribosomal subunit assembly. Proceedings of the National Academy of Sciences of the United States of America 112, E4707–4716, doi:10.1073/pnas.1506749112 (2015).

8 Connolly, K., Rife, J. P. & Culver, G. Mechanistic insight into the ribosome biogenesis functions of the ancient protein KsgA. Molecular microbiology 70, 1062–1075, doi:10.1111/j.1365-2958.2008.06485.x (2008).

9 Champney, S. Macromolecular Structure Assembly as a Novel Antibiotic Target. Antibiotics 11, doi:10.3390/antibiotics11070937 (2022).

10 Champney, W. S. Antibiotics targeting bacterial ribosomal subunit biogenesis. Journal of Antimicrobial Chemotherapy 75, 787–806, doi:10.1093/jac/dkz544 (2020).

11 Corrigan, R. M., Bellows, L. E., Wood, A. & Grundling, A. ppGpp negatively impacts ribosome assembly affecting growth and antimicrobial tolerance in Gram-positive bacteria. Proceedings of the National Academy of Sciences of the United States of America 113, E1710–1719, doi:10.1073/pnas.1522179113 (2016).

12 Zhang, Y. E. et al. (p)ppGpp Regulates a Bacterial Nucleosidase by an Allosteric Two-Domain Switch. Molecular cell 74, 1239–1249 e1234, doi:10.1016/j.molcel.2019.03.035 (2019).

13 Wang, B., Grant, R. A. & Laub, M. T. ppGpp Coordinates Nucleotide and Amino-Acid Synthesis in E. coli During Starvation. Molecular cell 80, 29–42.e10, doi:10.1016/j.molcel.2020.08.005 (2020).

14 Jeremia, L., Deprez, B. E., Dey, D., Conn, G. L. & Wuest, W. M. Ribosome-targeting antibiotics and resistance via ribosomal RNA methylation. RSC Medicinal Chemistry 14, 624–643, doi:10.1039/d2md00459c (2023).

15 Gutierrez, B. et al. Fitness Cost and Interference of Arm/Rmt Aminoglycoside Resistance with the RsmF Housekeeping Methyltransferases. Antimicrobial Agents and Chemotherapy 56, 2335–2341, doi:10.1128/aac.06066-11 (2012).

16 Wachino, J.-i., et al. Novel Plasmid-Mediated 16S rRNA m1A1408 Methyltransferase, NpmA, Found in a Clinically Isolated Escherichia coli Strain Resistant to Structurally Diverse Aminoglycosides. Antimicrobial Agents and Chemotherapy 51, 4401–4409, doi:10.1128/aac.00926-07 (2007).

17 Griffith, L. J., Ostrander, W. E., Mullins, C. G. & Beswick, D. E. Drug Antagonism between Lincomycin and Erythromycin. Science 147, 746–747, doi:10.1126/science.147.3659.746 (1965).

18 Greninger, A. L. et al. International Spread of Multidrug-Resistant Campylobacter coli in Men Who Have Sex With Men in Washington State and Québec, 2015–2018. Clinical Infectious Diseases 71, 1896–1904, doi:10.1093/cid/ciz1060 (2020).

19 Krüger, H. et al. Novel macrolide-lincosamide-streptogramin B resistance gene erm(54) in MRSA ST398 from Germany. Journal of Antimicrobial Chemotherapy 77, 2296–2298, doi:10.1093/jac/dkac149 (2022).

20 Schwarz, S., Werckenthin, C. & Kehrenberg, C. Identification of a Plasmid-Borne Chloramphenicol-Florfenicol Resistance Gene in Staphylococcus sciuri. Antimicrobial Agents and Chemotherapy 44, 2530–2533, doi:10.1128/aac.44.9.2530-2533.2000 (2000).

21 Yan, F. et al. RlmN and Cfr are Radical SAM Enzymes Involved in Methylation of Ribosomal RNA. Journal of the American Chemical Society 132, 3953–3964, doi:10.1021/ja910850y (2010).

22 Poteete, A. R. What makes the bacteriophage lambda Red system useful for genetic engineering: molecular mechanism and biological function. FEMS Microbiol. Lett. 201, 9–14, doi:10.1016/s0378-1097(01)00242-7 (2001).

23 Tu, S. et al. YcgC represents a new protein deacetylase family in prokaryotes. Elife 4, doi:10.7554/eLife.05322.00110.7554/eLife.05322.002 (2015).

24 Daan Frenkel & Smit., B. Understanding molecular simulation: from algorithms to applications. Vol. 1 (Elsevier, 2001).

25 Marrink, S. J. et al. Computational Modeling of Realistic Cell Membranes. Chemical Reviews 119, 6184–6226, doi:10.1021/acs.chemrev.8b00460 (2019).

26 Nagy, G., Gordillo, M. C., Guàrdia, E. & Martí, J. Liquid Water Confined in Carbon Nanochannels at High Temperatures. The Journal of Physical Chemistry B 111, 12524–12530, doi:10.1021/jp073193m (2007).

27 Calero, C., Martí, J. & Guàrdia, E. 1H Nuclear Spin Relaxation of Liquid Water from Molecular Dynamics Simulations. The Journal of Physical Chemistry B 119, 1966–1973, doi:10.1021/jp510013q (2015).

28 Jo;, S., Kim;, T., Iyer;, V. G. & Im, W. CHARMM-GUI: a web-based graphical user interface for CHARMM. J Comput Chem 29, 1859–1865, doi:10.1002/jcc.20945 (2008).

29 Kern, N. R., Lee, J., Kyo Choi, Y. & Im, W. CHARMM-GUI multicomponent assembler for modeling and simulation of complex multicomponent systems. Biophysical Journal 121, doi:10.1016/j.bpj.2021.11.2789 (2022).

30 Huang, J. & MacKerell, A. D. CHARMM36 all-atom additive protein force field: Validation based on comparison to NMR data. Journal of Computational Chemistry 34, 2135–2145, doi:10.1002/jcc.23354 (2013).

31 Berendsen, H. J. C., van der Spoel, D. & van Drunen, R. GROMACS: A message-passing parallel molecular dynamics implementation. Computer Physics Communications 91, 43–56, doi:10.1016/0010-4655(95)00042-e (1995).

32 Humphrey, W., Dalke, A. & Schulten, K. VMD: Visual molecular dynamics. Journal of Molecular Graphics 14, 33–38, doi:10.1016/0263-7855(96)00018-5 (1996).

33 Pettersen, E. F. et al. UCSF Chimera?A visualization system for exploratory research and analysis. Journal of Computational Chemistry 25, 1605–1612, doi:10.1002/jcc.20084 (2004).

34 Spangler, C., Bohm, A., Jenal, U., Seifert, R. & Kaever, V. A liquid chromatography-coupled tandem mass spectrometry method for quantitation of cyclic di-guanosine monophosphate. Journal of microbiological methods 81, 226–231, doi:10.1016/j.mimet.2010.03.020 (2010).

35 Purta, E., O’Connor, M., Bujnicki, J. M. & Douthwaite, S. YccW is the m5C Methyltransferase Specific for 23S rRNA Nucleotide 1962. J. Mol. Biol. 383, 641–651, doi:10.1016/j.jmb.2008.08.061 (2008).

36 Caldas, T. et al. The FtsJ/RrmJ Heat Shock Protein of Escherichia coli Is a 23 S Ribosomal RNA Methyltransferase. J. Biol. Chem. 275, 16414–16419, doi:10.1074/jbc.M001854200 (2000).

37 Ho, C. L. et al. Visualizing the perturbation of cellular cyclic di-GMP levels in bacterial cells. Journal of the American Chemical Society 135, 566–569, doi:10.1021/ja310497x (2013).

38 Lacanna, E., Bigosch, C., Kaever, V., Boehm, A. & Becker, A. Evidence for Escherichia coli Diguanylate Cyclase DgcZ Interlinking Surface Sensing and Adhesion via Multiple Regulatory Routes. Journal of bacteriology 198, 2524–2535, doi:10.1128/JB.00320-16 (2016).

39 Boehm, A. et al. Second messenger signalling governs Escherichia coli biofilm induction upon ribosomal stress. Molecular microbiology 72, 1500–1516, doi:10.1111/j.1365-2958.2009.06739.x (2009).

40 Chou, S. H. & Galperin, M. Y. Diversity of Cyclic Di-GMP-Binding Proteins and Mechanisms. Journal of bacteriology 198, 32–46, doi:10.1128/JB.00333-15 (2016).

41 Kramer, K. et al. Photo-cross-linking and high-resolution mass spectrometry for assignment of RNA-binding sites in RNA-binding proteins. Nature methods 11, 1064–1070, doi:10.1038/nmeth.3092 (2014).

42 Shu, C., Yi, G., Watts, T., Kao, C. C. & Li, P. Structure of STING bound to cyclic di-GMP reveals the mechanism of cyclic dinucleotide recognition by the immune system. Nature structural & molecular biology 19, 722–724, doi:10.1038/nsmb.2331 (2012).

43 Hou, Y.-J. et al. Structural insights into the mechanism of c-di-GMP–bound YcgR regulating flagellar motility in Escherichia coli. Journal of Biological Chemistry 295, 808–821, doi:10.1016/s0021-9258(17)49937-6 (2020).

44 Matsuyama, B. Y. et al. Mechanistic insights into c-di-GMP–dependent control of the biofilm regulator FleQ from Pseudomonas aeruginosa. Proceedings of the National Academy of Sciences 113, doi:10.1073/pnas.1523148113 (2015).

45 Yan, X.-F. et al. Structural analyses unravel the molecular mechanism of cyclic di-GMP regulation of bacterial chemotaxis via a PilZ adaptor protein. Journal of Biological Chemistry 293, 100–111, doi:10.1074/jbc.M117.815704 (2018).

46 Gupta, K., Liao, J., Petrova, O. E., Cherny, K. E. & Sauer, K. Elevated levels of the second messenger c-di-GMP contribute to antimicrobial resistance of Pseudomonas aeruginosa. Molecular microbiology 92, 488–506, doi:10.1111/mmi.12587 (2014).

47 Gomez, J. E. et al. Ribosomal mutations promote the evolution of antibiotic resistance in a multidrug environment. Elife 6, doi:10.7554/eLife.20420 (2017).

48 Hayes, C. S. et al. Adaptive Remodeling of the Bacterial Proteome by Specific Ribosomal Modification Regulates Pseudomonas Infection and Niche Colonisation. PLoS genetics 12, doi:10.1371/journal.pgen.1005837 (2016).

49 Søgaard-Andersen, L. et al. Control of mRNA translation by dynamic ribosome modification. PLoS genetics 16, doi:10.1371/journal.pgen.1008837 (2020).

50 Guo, Q. et al. Elongation factor P modulates Acinetobacter baumannii physiology and virulence as a cyclic dimeric guanosine monophosphate effector. Proceedings of the National Academy of Sciences 119, doi:10.1073/pnas.2209838119 (2022).

51 Inuzuka, S. et al. Recognition of cyclic-di-GMP by a riboswitch conducts translational repression through masking the ribosome-binding site distant from the aptamer domain. Genes to Cells 23, 435–447, doi:10.1111/gtc.12586 (2018).

52 Davis, J. H. & Williamson, J. R. Structure and dynamics of bacterial ribosome biogenesis. Philosophical Transactions of the Royal Society B: Biological Sciences 372, doi:10.1098/rstb.2016.0181 (2017).

53 Pletnev, P. et al. Comprehensive Functional Analysis of Escherichia coli Ribosomal RNA Methyltransferases. Frontiers in Genetics 11, doi:10.3389/fgene.2020.00097 (2020).

54 Osterman, I. A., Dontsova, O. A. & Sergiev, P. V. rRNA Methylation and Antibiotic Resistance. Biochemistry (Moscow) 85, 1335–1349, doi:10.1134/s000629792011005x (2020).

55 Blair, J. M., Webber, M. A., Baylay, A. J., Ogbolu, D. O. & Piddock, L. J. Molecular mechanisms of antibiotic resistance. Nature reviews. Microbiology 13, 42–51, doi:10.1038/nrmicro3380 (2015).

56 Darby, E. M. et al. Molecular mechanisms of antibiotic resistance revisited. Nature Reviews Microbiology 21, 280–295, doi:10.1038/s41579-022-00820-y (2022).

